# Spectral Contrast Optical Coherence Tomography Angiography Enables Single-Scan Vessel Imaging

**DOI:** 10.1101/406652

**Authors:** James A. Winkelmann, Aya Eid, Graham Spicer, Luay M. Almassalha, The-Quyen Nguyen, Vadim Backman

## Abstract

Optical coherence tomography angiography relies on motion for contrast and requires at least two data acquisitions per pointwise scanning location. We present a method termed spectral contrast optical coherence tomography angiography using visible light that relies on the spectral signatures of blood for angiography from a single scan using endogenous contrast. We demonstrate the molecular sensitivity of this method, which enables lymphatic vessel, blood, and tissue discrimination.

## Main Text

Optical coherence tomography (OCT) is a noninvasive optical imaging modality that provides micron-scale resolution of three-dimensional (3D) tissue morphology^1^. In addition to providing structural information, enhanced processing of an OCT signal can provide a milieu of functional and molecular information^2–4^. Early work harnessed the Doppler phase shift caused by backscattering from erythrocyte motion within vessels to measure blood flow velocity and delineate vessels, called Doppler OCT^5,6^. To achieve this motion-based contrast enhancement, algorithms utilizing phase variance, sequential scan subtraction, and speckle variance, among others, were developed and successfully implemented to enhance vasculature in OCT scans^7–11^. The limitations of motion-based OCT angiography include sensitivity to breathing and pulse movement in living animals, which commonly result in bright banding artifacts across the projected OCT angiogram^3,12^.

In addition to motion-based OCT angiography, spectroscopic visible band OCT imaging has enabled true-color imaging of biological tissues by resolving distinct spectral absorption features^2^. The ability to quantify hemoglobin concentration and oxygenation from endogenous contrast and molecular information from exogenous nanoparticle-based contrast agents are promising applications made possible through the development of visible spectroscopic OCT (Supplementary Note 1)^13–20^. Thus, the development of OCT systems in the visible region has provided a promising avenue for the measurement of valuable absorption-based information at high spatial resolution.

Herein, we present a novel, facile and robust method of obtaining angiography images from a spectral domain OCT (SD-OCT) signal, called spectral contrast OCT angiography (SC-OCTA). Utilizing distinct spectral features of hemoglobin in the visible range, SC-OCTA enables 3D angiography without the need to repeat scanning protocols, eliminating all motion-based artifacts ubiquitous in previously established OCTA and allowing for the fastest SD-OCT angiography acquisition speeds to date. Furthermore, this unique method of spectral-based vessel segmentation eliminates the need for blood-flow-induced motion for angiography, allowing for the novel ability to image vasculature in hemostatic tissues, such as hemorrhage from compromised vasculature in the case of cardiovascular disease^21^.

While there are several OCT operating regions that can be selected, a typically favored range for biological imaging is the near-infrared (NIR) region from approximately 700–900 nm. This is because the short wavelength range provides a higher OCT axial resolution than at wavelengths greater than 1000 nm, and the region falls within the ‘optical window.’ In the ‘optical window,’ there is minimal absorption from water and hemoglobin, allowing for high penetration in tissue. This allows NIR OCT systems to penetrate deeply into tissue but diminishes sensitivity to blood and tissue scattering spectral features. Blood absorption coefficients are two orders of magnitude higher in the 400–600 nm range, and the tissue scattering coefficients are approximately double compared with the those in the NIR range^22^, allowing visible OCT systems to be sensitive to blood oxygenation and achieve a higher image contrast^2,23^.

In visible and NIR spectra, tissue exhibits a monotonic decrease in scattering with increasing wavelength. This relationship follows a power-law behavior that is directly related to the fractal nature of tissue when modeled as a continuous random medium^4^. However, the scattering spectrum from hemoglobin follows nonmonotonic behavior; between 550 and 600 nm, hemoglobin scattering increases with increasing wavelength^22,24^. We exploit this unique spectral feature, combined with high image contrast from the visible spectrum, to rapidly and easily image tissue and blood with a clear discrimination of vessels down to the level of individual capillaries.

SD-OCT obtains depth-resolved sample information by taking a Fourier transform of the interference recorded as a function of wavelength (on a spectrometer) between a reference reflection and light scattered from the sample (Figure 1 (a)). By subsampling the spectrum with a short time Fourier transform (STFT), spectrally dependent OCT A-lines are measured. Therefore, opposite spectral slopes of blood and tissue can be spatially visualized by looking at the contrast of spectrally dependent OCT image intensities from 550 nm to 600 nm. We found that a Kaiser sampling window at 557 nm and 620 nm with a full width at half maximum (FWHM) of ~38 nm provided high spectral contrast between blood and the surrounding tissue.

**Figure 1.**
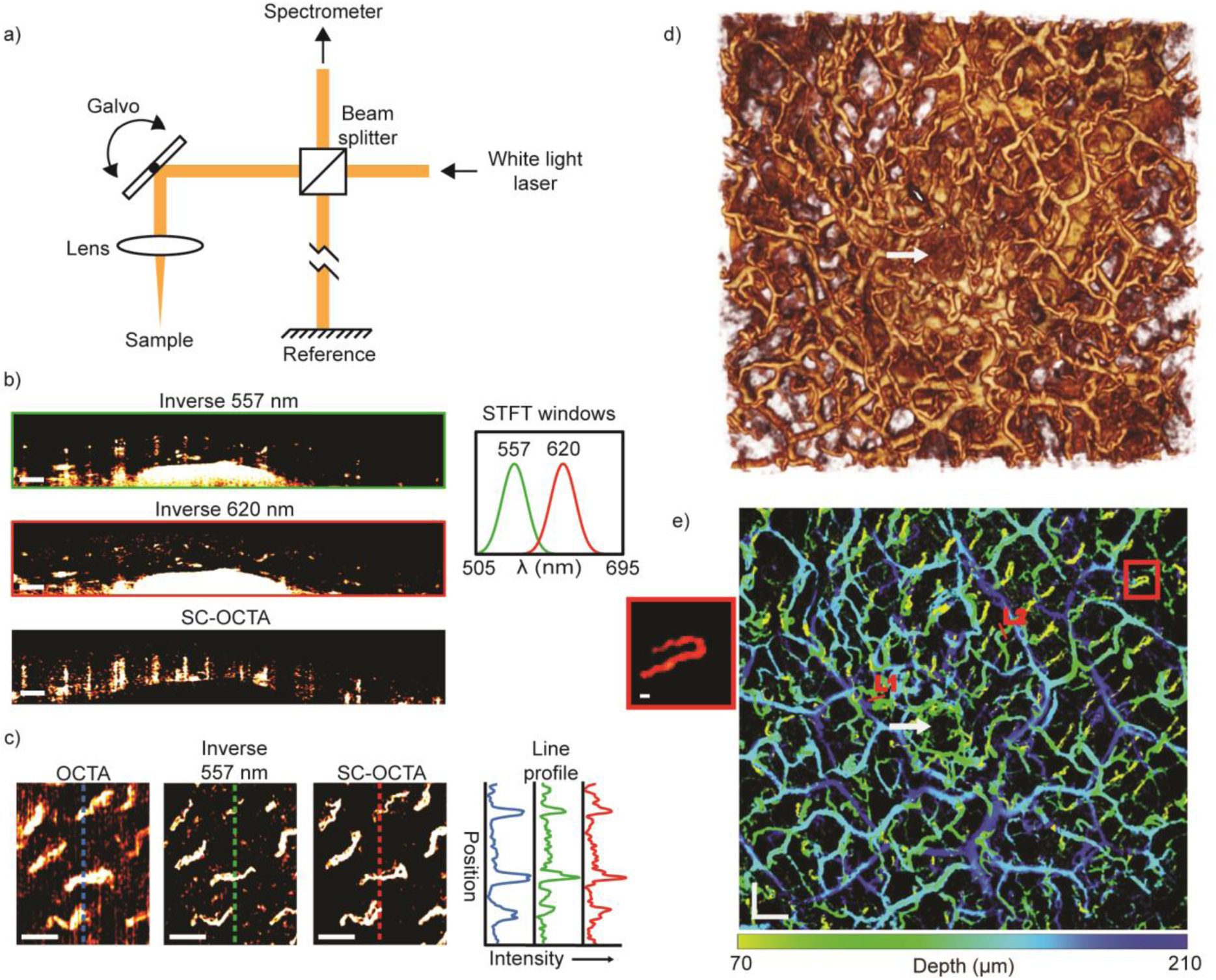
(a) Simplified schematic of the visible OCT system, which allowed 3D spectral information of the sample to be obtained. (b)-(e) *In vivo* human imaging of labial mucosa (lower lip) from a healthy volunteer. (b) Inverse 557 nm and inverse 620 nm B-scans with their corresponding STFT windows, as well as a SC-OCTA B-scan showing contrast shadows from each vessel. Scale bar: 200 µm. (c) Comparison of angiography *en face* projections of superficial capillary loops with traditional motion contrast OCTA (64–111 µm), inverse 557 nm (55.6–140 µm), and SC-OCTA (83–209 µm) with their corresponding line profile intensities. Depth ranges were chosen to maximize the contrast of the *en face* projections for different techniques. Scale bar: 200 µm. (d) 3D rendering of inverse 557 nm. Scan area: 3.65 × 3.44 mm. (e) Depth-encoded vessel map of the same FOV as (d) with saturation and value from SC-OCTA and hue from the depth of the vessel in inverse 557 nm. Line 1 (L1) and Line 2 (L2) are line profiles in Figure S3 for comparison with the simulated line profile results. Scale bars: 300 µm. The red box shows a blow-up of the capillary loop; scale bar 20 µm. (d & e) The white arrow shows a salivary duct that is correctly not identified by SC-OCTA in (e).

*In vivo* B-scans (Figure 1 (b)) of lower human labial mucosa (inner side of lip) can be seen with the ratio of the OCT image intensities from the two Kaiser windows (620 nm divided by 557 nm), hereafter referred to as SC-OCTA. In images from the inverse OCT intensity at 557 nm, hereafter referred to as inverse 557 nm images, blood vessels can easily be seen due to the high contrast and high absorption provided in the visible range. The spectral contrast image demonstrates how blood vessels are highlighted by a shadow and tissue is ignored. For the OCT system (Figure S1), the experimental air axial resolution of these Kaiser windows was found to be 3.80 µm and 4.72 µm for the 557 nm window and 620 nm window, respectively (Figure S2). According to Beer’s law and Mie theory simulations, the SC-OCTA method should work all the way down to ~4 µm diameter capillaries with only a single scan (Supplementary Note 2). To confirm capillary imaging, inverse 557 nm and SC-OCTA *en face* projections were compared with the results of traditional OCTA phase and amplitude contrast (Figure 1 (c)). The same eight capillary loops in the labial mucosa are seen in the inverse 557 nm image, SC-OCTA, as well as in the traditional OCT angiography, which requires the sample to be scanned at least twice. It took 18.2 sec to acquire the traditional OCTA data and an effective 4.5 sec for SC-OCTA data. Fourier ring correlation analysis was performed using a half-bit threshold computed an effective resolution of 20.19 µm for the traditional OCTA, 12.2 µm for inverse 557 nm and 8.92 µm for the SC-OCTA method (Figure S4)^25^. This analysis demonstrates how inverse 557 nm and SC-OCTA are less sensitive to *in vivo* motion than OCTA. A detailed large field of view of the labial mucosa (Figure 1 (d & e)) demonstrates the ability of SC-OCTA to resolve arteriolar and capillary-level vessels (Figure S5) with only a single A-line acquired at each point-scanning location. The inverse 557 nm image does not differentiate low scattering structures from hemoglobin absorption. This is noted by the white arrow showing a salivary duct that is visible in the inverse 557 nm image (Figure 1 (d)) but not in the SC-OCTA image (Figure 1 (e)).

Because SC-OCTA does not rely on motion for contrast, it can image nonflowing blood and highly moving samples. To demonstrate this capability, we fabricated an ~55 µm diameter vessel phantom (Figure S6 (a & b)) and recorded the signal-to-noise ratio (SNR) of the SC-OCTA and OCTA signals under different flow conditions (Figure 2) and vibrations (Figure S6 (d & e)). The results showed that in contrast to OCTA, the SC-OCTA signal is not significantly affected by flow and can image highly moving samples. With a limited vessel phantom lifetime, waiting for blood turbulence to approach zero was not possible. Therefore, to demonstrate the utility of SC-OCTA in the hemostasis setting, the serosal surface of a freshly sacrificed mouse large intestine was imaged (Figure 3). To the best of our knowledge, this is the first time angiography has been performed on tissue with nonmoving blood with endogenous contrast using OCT. The results show that OCTA struggles to resolve any vessels in the hemostasis setting, while SC-OCTA can detect several vessels with a faster acquisition time (Figure 3 (a)). To demonstrate the molecular sensitivity of SC-OCTA, lymphatics and blood vessels were imaged on a freshly sacrificed mouse omentum (Figure 4 (a-d)) and a heart surface, where images were compared with histological imaging of the same tissue (Figure 4 (e-g)). Notably, SC-OCTA differentiates blood vessels from low-scattering lymphatic vessels and adipocytes. A B-scan of depth-integrated SC-OCTA, where each pixel in the SC-OCTA image is integrated 50 µm along the depth and multiplied by the inverse 557 nm image, allowed vessels to be represented in three dimensions (Figure 4). The depth integration technique allowed labial mucosa vasculature to be differentiated from the salivary duct and tissue in three dimensions all the way down to the capillary level (Video S1). Depth-integrated SC-OCTA demonstrated the ability to image the branches of the coronary arteries and differentiate these from neighboring lymphatic vessels (Figure 4 (e), Video S2). The high resolution and contrast in the visible spectrum additionally allowed imaging of lymphatic valves where the tricuspid structure of the valve is easily discerned in three dimensions (Figure 4 (d), Video S3).

**Figure 2.**
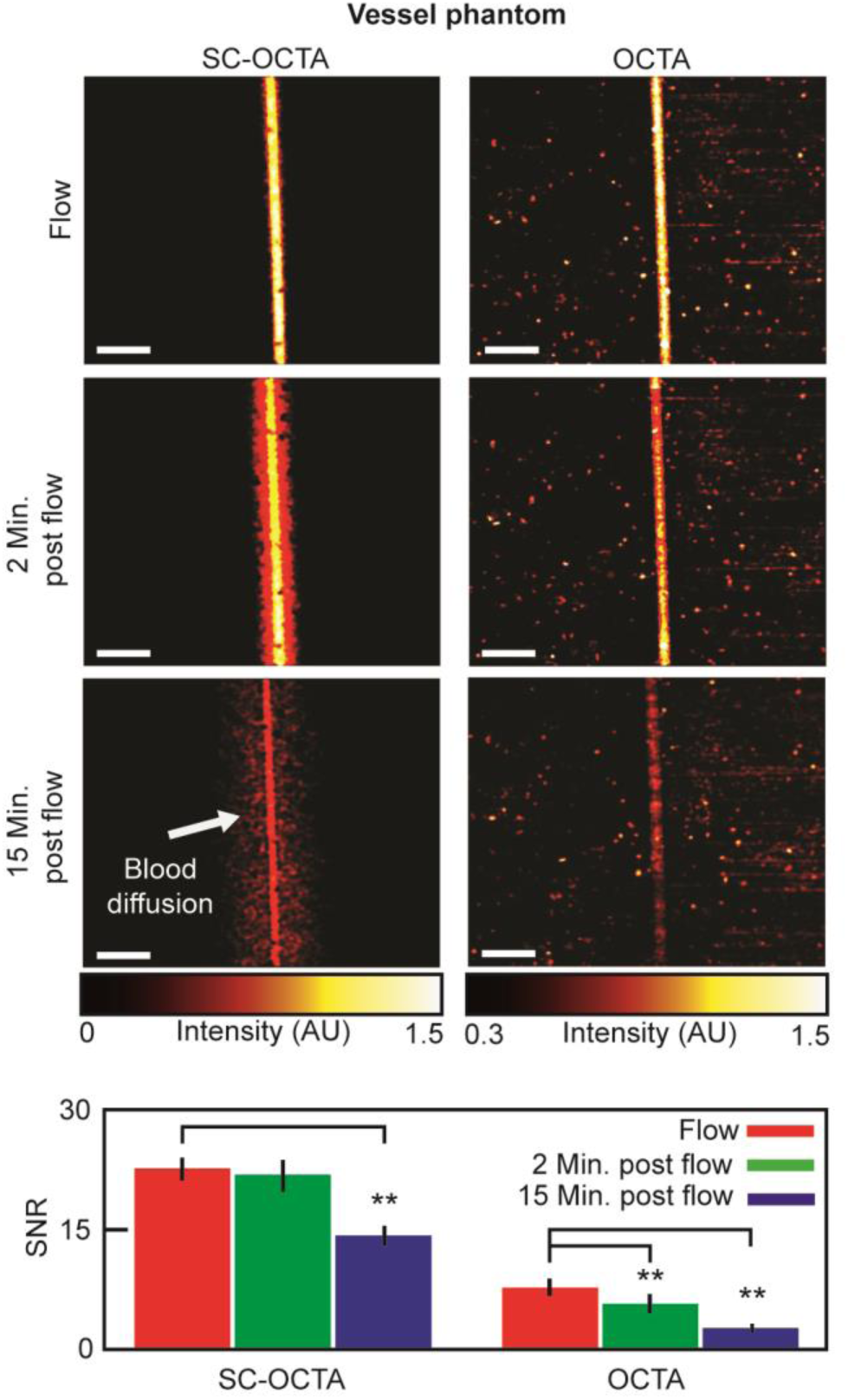
Vessel phantom imaging using bovine blood. *En face* projections of SC-OCTA and OCTA under different flow conditions and the corresponding signal-to-noise ratio (SNR). Due to the dynamic nature of the agarose-based phantom, SC-OCTA measurements were taken from one of the repetitions of the OCTA scan to ensure similar measurement conditions. SC-OCTA (flow: 22.76±1.42; 2 min post flow: 21.99±1.95; 15 min post flow: 14.08±1.30). OCTA (flow: 7.69±1.13; 2 min post flow: 5.74±1.21; 15 min post flow: 2.35±0.60). ** (p<.01) for the two-sample t-test. The standard deviation is taken over 10 equally sized regions across the phantom (Figure S6 (f)). Scale bar: 250 µm. During the flow measurement, a syringe pump supplying blood to the phantom was set to .0006 µL/sec. Measurements were then taken 2 min and 15 min after stopping the flow to the phantom. It can be seen that SC-OCTA SNR is not significantly affected 2 min after stopping the flow, and SC-OCTA can visualize blood diffusing into the vessel phantom (Figure S6 (a)) not seen by OCTA. The SNR is significantly affected 15 min post flow in SC-OCTA due to decrease in the blood concentration in the vessel channel. The SNR was computed from the raw *en face* projection intensities, while the images of the *en face* projections were scaled to minimize their background intensities.

**Figure 3.**
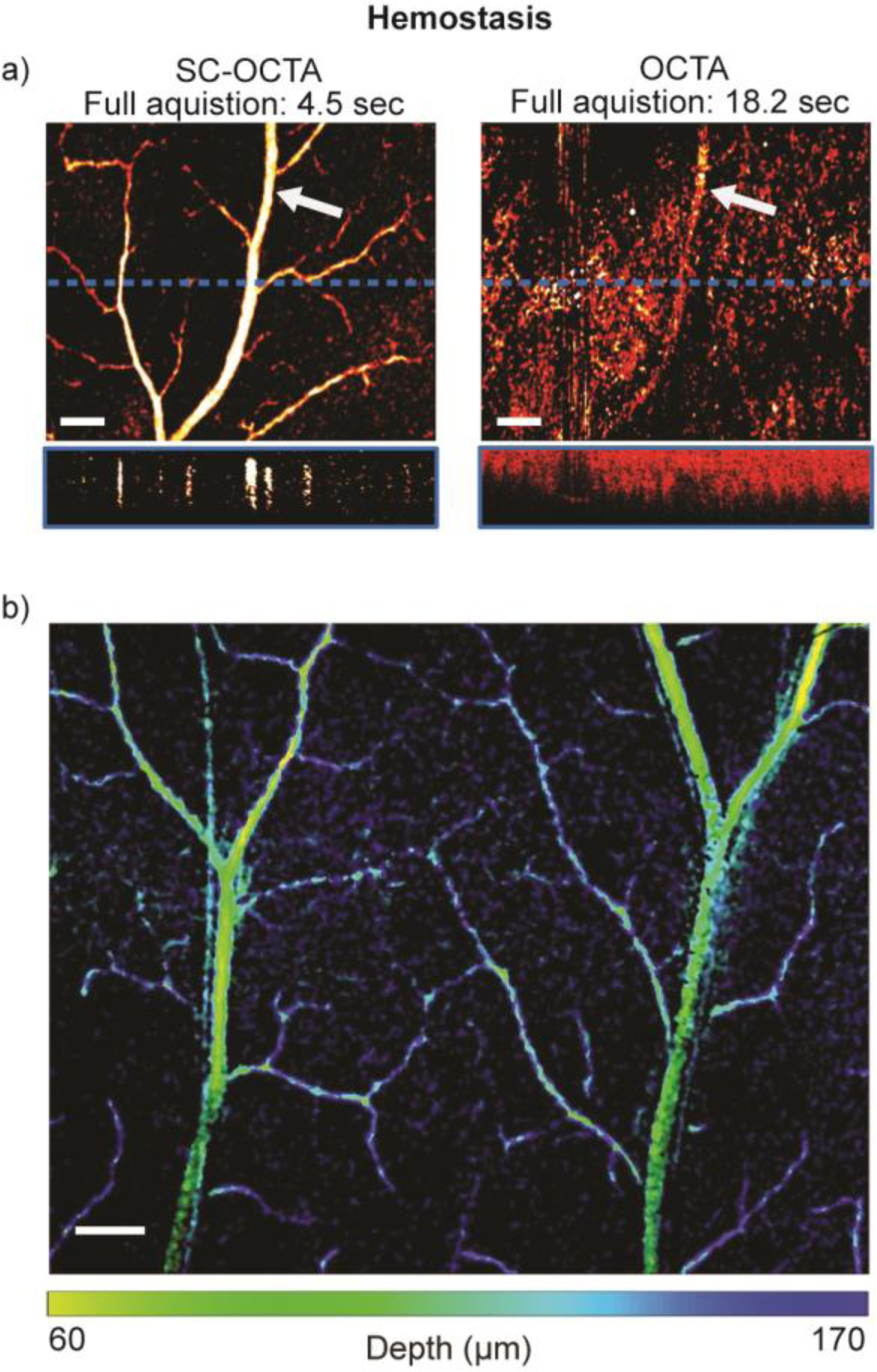
Sacrificed mouse hemostasis imaging of serosal surface of large intestines. (a) Comparison of SC-OCTA and OCTA *en face* projections and B-scans (from the blue dotted line on the *en face* projection). The power of SC-OCTA can be visualized in the case of hemostasis, as OCTA has difficulty sensing even large vessels. The white arrow points to the same vessel detected by SC-OCTA (56–280 µm) and OCTA (28–336 µm). The depths were chosen to maximize the contrast of each method. Scale bar: 200 µm. (b) Large field of view from SC-OCTA with saturation and value from SC-OCTA and hue from the depth of vessel in the inverse 557 nm. Scale bar: 250 µm. Please note this is serosal surface imaging, and the low-signal capillary loops, located just beneath the surface from the luminal side of the large intestine, were not clearly visualized by SC-OCTA^29^.

**Figure 4.**
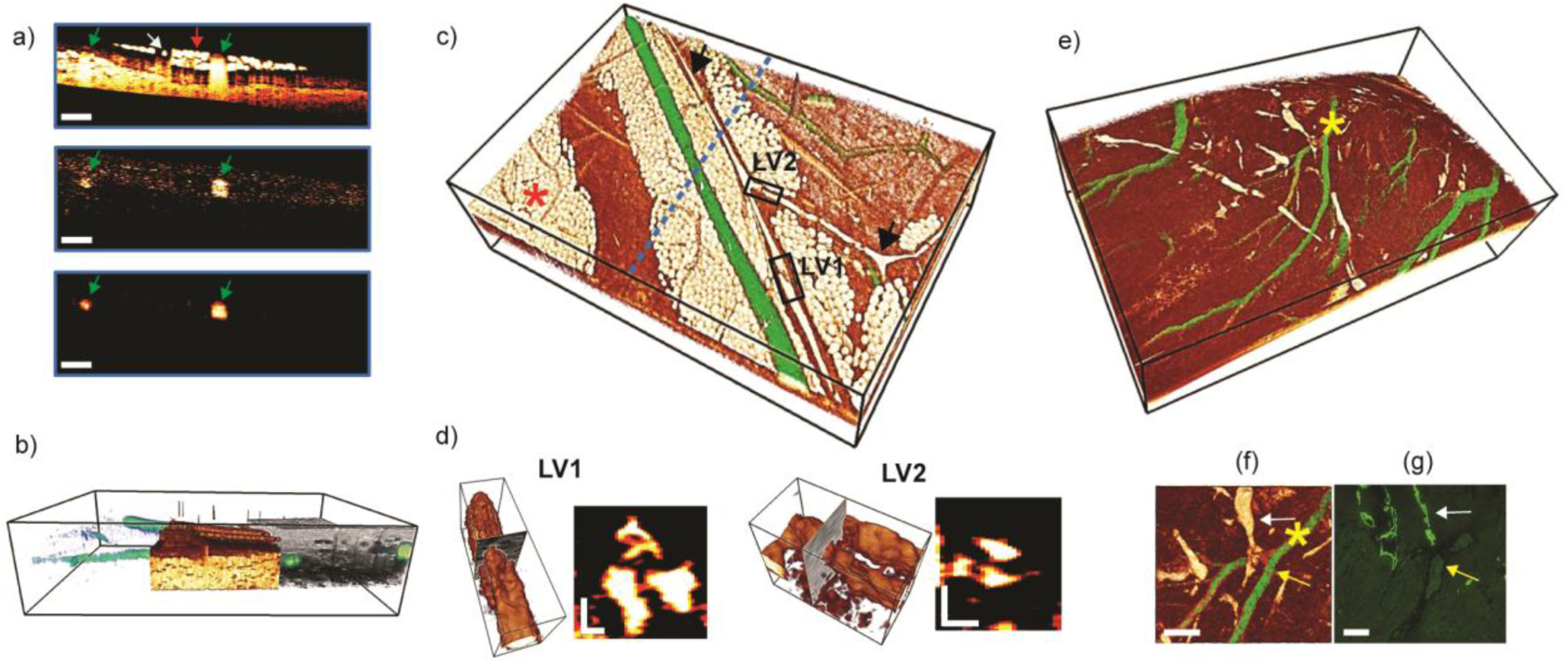
Single-scan vessel imaging of blood and lymphatic vessels of a sacrificed mouse anterior abdominal wall (a-d) and heart surface (e-f). (a) B-scan comparison of inverse 557 nm, SC-OCTA, and depth-integrated SC-OCTA showing how blood is only highlighted using spectral contrast. Blood vessels (green arrow), lymphatic vessel (white arrow), and adipocytes (red arrow). Scale bar: 250 µm. (b) Side view peel-away showing depth-integrated SC-OCTA, which allowed blood vessels to be visualized (green), inverse 557 nm with vessels removed showing adipose/lymphatic tissue (White/Orange) and full-spectrum 505-695 nm OCT intensity showing highly scattering tissue (gray). Bounding box: 2.52 × 3.78 × 0.7 mm. (c) Color-coded 3D rendering. Depth-integrated SC-OCTA (green) showing blood vessels and inverse 557 nm (White/Orange) showing adipocytes (red asterisk) and lymphatics (black arrows). The blue dotted line shows the cross-section location of (a). Bounding box: 2.52 × 3.78 × 0.7 mm. (d) 3D rendering and B-scan cross-sections of lymphatic valve 1 (LV1) and lymphatic valve 2 (LV2) from the black boxes in (c), showing a tricuspid valvular structure. Bounding box for LV1: 336 × 112 × 105 µm. Bounding box for LV2: 256 × 141 × 130 µm. Scale bars: 30 µm. (e) Color-coded 3D rendering. Depth-integrated SC-OCTA (green) showing blood vessels and inverse 557 nm (White/Orange) showing lymphatics in white. Bounding box: 2.02 × 3.36 × 1 mm. (f) Top view of (e) showing a blood vessel branch (yellow arrow) with lymphatic vessels (white arrow) and (g) corresponding immunofluorescence microscopy localizing podoplanin to distinguish blood vessels from lymphatic vessels. (f-g) Scale bars: 50 µm. The yellow asterisk in (e) corresponds to the yellow asterisk in (f).

In summary, we demonstrate a robust method for single-scan angiography and tissue differentiation with molecular sensitivity in three dimensions using spectroscopic visible OCT (Video S4, Figure S7). Future plans will aim to further optimize SC-OCTA algorithms and implement visible OCT endoscopy for minimally invasive *in vivo* imaging with molecular sensitivity^26^.

## Methods

Please see Figure S8 for a flow chart of the spectral-contrast-based angiography (inverse 557 nm, SC-OCTA, depth-integrated SC-OCTA) processing steps.

### SD-OCT setup

The following OCT setup description is in reference to the schematic shown in Figure S1. A supercontinuum laser (NKT Photonics, SuperK Extreme EXW-6) was set to 100% output power, and the direct output from the laser was first sent through a splitter box (NKT Photonics, SpectraK Split) that contained a 400–850 nm reflective bandpass filter to dump the infrared parts of the spectrum. This splitter box is not shown in Figure S1. The spectrum of light was then smoothed using two prisms and a spatial filter to achieve a similar dynamic range across the full spectrum. Light first passed over a pickoff mirror and into Prism 1 (Thorlabs, F2 equilateral prism, PS854). The incident angle of the beam onto Prism 1 was set to the minimum angle of deviation to minimize reflections off prism-air interfaces. Prism 1 refracted the light and angularly dispersed the light as a function of wavelength. After a sufficient distance to spread out the spectrum in space, the light entered Prism 2 (Thorlabs, F2 equilateral prism, PS858). Prism 2 was adjusted so that the incident surface was parallel to the output surface of Prism 1. Prism 2 recollimated the light but with the beam being dispersed in wavelength across its horizontal axis. A piece of matte black aluminum foil (Thorlabs, BKF12) was cut to the shape of an oval and attached to a 2-dimensional translational mount to act as a spatial filter. The translational mount allowed for fine tuning of the spectrum as the foil attenuated part of the beam cross-section. The light was then reflected off a mirror that slightly deviated the beam downwards to allow the returning beam through the prisms to be reflected by the pickoff mirror. The light was then passed through a linear polarizer (Newport, 10LP-VIS-B) and coupled into 7 meters of SM 600 fiber (Thorlabs, 900 µm tight buffer) with Objective 1 (Edmund Optics, 33–438). The SM 600 fiber was threaded through two sets of three-paddle polarization controllers (Thorlabs, FPC562), where only 2 paddles were used on one of the controllers (three paddles were used in one controller and two were used in the second). A linear polarizer and two sets of three-paddle polarization controllers were necessary to have sufficient polarization control to maximize the interference efficiency of the OCT interferometer across its broad bandwidth. Light was collimated out of the SM600 fiber using a fiber port collimator (OZ Optics, HPUCO-23–400/700-S-10AC) to a cube 50:50 beam splitter (Thorlabs, CM1-BS1), which directed light to the sample arm and reference arm. In the sample arm, a two-dimensional mirror galvanometer (Thorlabs, GVS002 TSH25379-X) allowed the beam to be pointwise scanned across the sample. The beam was focused on the sample using Objective 2 (Thorlabs, LSM03-VIS). The reference arm contained a dispersion compensator (Thorlabs, LSM03DC-VIS), which was specifically made for Objective 2. A razor blade was used to attenuate the beam in the reference arm so that the reference power was within the dynamic range of the spectrometer. The reference mirror in the reference arm was placed on a translation stage to allow for fine adjustment of the reference arm path length with respect to the sample arm path length. A fiber port collimator (OZ Optics, HPUCO-23–400/700-S-10AC) collected the interfered beam into SM-460B fiber (Thorlabs, P1–460B-FC-5), which directed the light to a custom-built visible spectrometer. Light was focused onto a 1200 lines/mm grating (Wasatch Photonics) from the SM-460B fiber with a mirror fiber collimator (Thorlabs, RC12APC-P01). The grating angularly dispersed the light as a function of wavelength onto a custom built 6-element focusing objective (effective focal length = 123.7 mm). The custom objective focused the light onto a 4096 × 2 pixel line scan camera (Basler, spL4096–140km). The mirror collimator, grating, and custom objective were placed on a translational mount to allow fine tuning of the distance between the components and the line scan camera. A custom spectrometer across such a broad bandwidth can be particularly challenging to construct and align.

### System Sensitivity and Resolution

Using the impulse response of a mirror, the sensitivity of the system was found to be 91.61 dB at an illumination power of 11.2 mW. The mirror impulse response was also used in calculating the air axial resolution, which was found to be 1.53 µm, corresponding to a tissue axial resolution of ~1.15 µm. The air axial resolution for the two Kaiser windows used in SC-OCTA was found to be 3.8 µm and 4.72 µm for the 557 nm window and 620 nm window, respectively. The air axial imaging range for the system was 1 mm with a roll-off sensitivity of approximately −10 dB/mm. The system sensitivity measurements can be seen in Figure S2. The lateral resolution was found to be 8.11 µm by measuring the edge response of a razor blade placed at the focal point. The first spatial derivative of the OCT intensity across the razor blade was computed using a Savitzky-Golay filter, and its full width at half-maximum was computed to give the lateral spatial resolution of the system. This same method was used to determine the resolution of the high numerical aperture system setup used in obtaining Figure S5 (c).

### Acquisition Parameters

The spectrometer for all data collected in this paper was set to 45,000 A-lines/sec at an exposure time of 18 µsec. Data collected for OCTA consisted of 4 repetitive B-scans containing 400 A-lines at each cross-section and 512 B-scans in the C-scan direction, taking a total of 18.2 sec to cover the full 1.76 × 1.76 mm field of view. To ensure similar measurement conditions, OCT data collected for the inverse 557 nm and SC-OCTA images in Figure 1 (c) and Figure 2 were collected from only 1 B-scan repetition of the OCTA scanning protocol, resulting in an effective acquisition time of 4.5 sec. The SC-OCTA phantom vibration data in Figure S6 (c & e) and the imaging of hemostasis in Figure 3 (a) were acquired in 4.5 sec. This is because imaging the phantom as quickly as possible can help minimize the motion between B-scans, which is important to ensure proper median filtering during SC-OCTA processing. For the hemostasis imaging case, we did not expect there to be a large temporally varying component in the minute between SC-OCTA single repetition acquisition and OCTA multirepetition acquisition. All other data collected in this paper were acquired by scanning a field of view of 3.78 × 3.78 mm and contained 900 × 900 A-lines, taking 18 sec.

### Axial PSF and System Roll-off Calibration

Spectroscopic OCT data were normalized by an aqueous calibration solution, which was measured following sample imaging. It is important to calibrate immediately after an imaging session because changes in polarization or reference arm position can change calibration data. In a perfectly static system, only one calibration should be required, but if the fibers in the system have slightly moved between imaging sessions, this can affect polarization and change the interference efficiency across the spectrum, leading to alterations in the relative intensity of the sampling windows. Likewise, if the reference arm position changes with respect to the focal point of the objective, the intensity of the sampling windows will be altered as a function of depth. The aqueous solution consisted of 80 nm sulfate latex beads (Molecular Probes by Life Technologies, 8% w/v) diluted to a concentration of 1% in deionized water. The solution was placed on a piece of angled quartz glass and imaged at 9 equally spaced locations in the axial direction using a 3D stage (Zaber, X-XYZ-LSQ150B-K0060-SQ3). The starting bead surface location was ~150 µm from the reference-sample zero-path length difference, and the ending bead surface position was ~950 µm from the reference-sample zero-path length difference. The OCT intensity for each STFT window from 1.4 µm to 8.4 µm into the bead solution was averaged for each depth location and then interpolated along the depth to obtain an axial intensity calibration for each STFT window.

### Raw Interferogram Data Processing

Interferogram data (data collected from the spectrometer) were processed in MATLAB utilizing a CPU and GPU. The raw interferogram data first had their direct current component removed and then was normalized to the reference arm intensity. The data were then multiplied by the sampling window so that an STFT could be performed. Kaiser sampling windows were chosen for spectral-contrast-based angiography to reduce sidelobes and reduce the transition band. Dispersion correction was also applied when necessary^27^. The data were then interpolated to be equally spaced in wavenumber space and fast Fourier transformed on the GPU. The data were then divided by the axial calibration intensity, squared, and multiplied by the center wavenumber of the sampling window raised to the fourth power. To summarize, the spectrally dependent OCT A-line intensity, *I*(*k*,*z*), was calculated using the following:

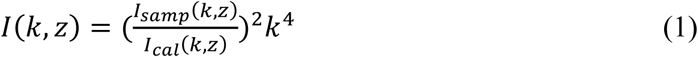
where *k* is the wavenumber (2π/wavelength), *z* is the depth along the A-line, and *I_samp_*(*k,z*) and *I_cal_*(*k,z*) are the STFTs of the sample and axial intensity calibration, respectively.

### Edge Detection

The surface of the sample needed to be calculated for computing the depth of blood vessels and removing the air on top of the sample for the inverse 557 images. The upper surface of the sample was determined by a series of morphological operations on each B-scan. This included smoothing each scan using Gaussian and median filters, contrast enhancing, and applying an extended maximum transform to find the largest continuous region of high contrast scattering. The parameters of each operation were heuristically determined for each sample. The surface points were calculated for each B-scan, and the 2D surface map was filled using a surface extrapolate and smoothed.

### OCTA Processing

The OCTA *en face* projections were generated using a phase sensitive decorrelation algorithm^17^. The OCT data from consecutive B-scans were first corrected for global phase fluctuations using a phase modifier in the axial and B-scan directions. The difference between the second to fourth consecutive B-scans was then calculated. This was performed for STFT Gaussian windows centered at 593.96 nm, 615.54 nm, and 638.74 nm, all with a FWHM of ~50 nm. The OCTA data produced for all the Gaussian windows and subtractions were then averaged to produce the final 3D OCTA data for the *en face* projection image. STFTs were used to reduce the OCTA axial resolution and make the method less sensitive to sample bulk motion^28^. It has also been shown in a previous study that this reduction in axial resolution should not hinder the ability of OCTA to sense capillary level vessels^20^.

### Inverse 557 nm OCT Intensity Processing

Inverse 557 nm OCT intensity data, 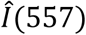, were produced by the following:

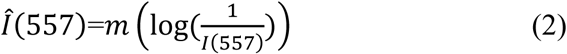
where *I*(557) is the 3D spectrally dependent OCT data of the 557 nm Kaiser window and *m* denotes a 3 A-line X 3 A-line X 4.2 µm (B-scan direction, C-scan direction, depth direction) median filter. The air surface above the sample was removed using the aforementioned edge detection algorithm. The 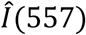 data shown for the labial mucosa in Figure 1 (d) had additional processing steps, including connected component analysis followed by a binary opening operation that was multiplied by the original 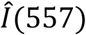 data. The rest of the 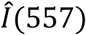 data shown in this brief communication did not have connected component analysis or opening operations.

### SC-OCTA Processing

The OCT data for each Kaiser window were dispersion compensated or axially shifted to coregister the two windows. This was a crucial step to ensure that edges were not highlighted in SC-OCTA due to poor coregistration. The 3D SC-OCTA intensity, *I_SC-OCTA_*, was calculated using the following:

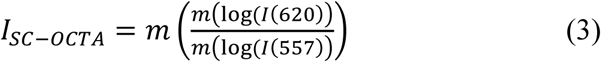
where *m* is the same size as that used in (2) and *I*(620) is the 3D spectrally dependent OCT data of the 620 nm Kaiser window.

The 3D depth-integrated SC-OCTA, *I_DI,SC-OCTA_*, was calculated using the following:

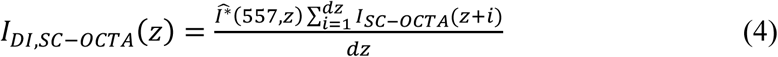
where 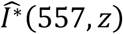 is 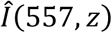 computed in (2) rescaled between 0 and 1, and *dz* is the depth integration amount.

### Blood, Tissue, Lymphatic, and Fat Region Backscattering Spectra Calculation

To extract the normalized backscattering spectra, μ*_b_*(*k*), of each tissue type (Figure S7), 3D masks were created to isolate each of the following: vascular and lymphatic networks, adipocytes, and tissue. The mask for the vascular network was generated using a simple binary threshold on the depth-integrated SC-OCTA image. Morphological operations, including binary opening and eroding, were applied with an effect size chosen heuristically to ensure that all voxels in the mask were safely within the blood vessel domains. The lymphatic network was manually segmented from the inverse 557 nm image. Adipose cells were segmented using an extended minimum transform on the full-spectrum OCT intensity image to find large continuous blobs of low-scattering regions; however, the cells in the axial path of blood vessels were avoided. The tissue was extracted based on a threshold range while avoiding the above expanded masks.

Each 3D mask was applied to a hyperspectral cube consisting of 34 wavenumbers, and the median spectra were computed and plotted in Figure S7. A 34-window spectral cube was generated with a STFT using a Gaussian window with a FWHM of 0.37 µm^-1^; the windows were linearly spaced in wavenumber. The normalized μ*_b_*(*k*) was related to *I*(*k*) utilizing the relation, μ*_b_*(*k*)~*I*(*k*).

### Vessel Phantom

To mimic tissue, a vessel phantom consisting of water, 4% agarose (Fischer Bioreagents), and 1% aqueous 80 nm polystyrene beads (Molecular Probes by Life Technologies, 8% w/v) was prepared (Figure S6 (a)). A 3D printed mold was made to hold the phantom. First, FEP tubing (outer diameter: 800 µm, inner diameter 250 µm; The Dolomite Center Ltd) was threaded into the 3D printed mold to serve as a conduit to deliver blood to the phantom. From the opposite side of the mold, 50 µm diameter tungsten wire (Malin Co.) was threaded into the mold and into the opening of the FEP tubing. A 3D printed spacer was then placed on top of the mold to create an ~100 µm gap between the spacer and the tungsten wire. Agarose and water were mixed and heated. Once the agarose was dissolved, an aqueous 80 nm polystyrene bead solution was added, and the mixture was poured into the mold. After the solution solidified in the mold, the spacer was carefully removed, and the tungsten wire was pulled out, creating an ~45 µm diameter channel that expanded to ~55 µm after hepranized bovine blood (Quad Five) was flown through at a rate of 0.0006 µL/sec with a syringe pump (Harvard Apparatus PhD 2000) (Figure S6 (b)). Since the phantom was made of agarose, this limited its operation lifetime due to blood diffusion; therefore, a new phantom was made for each experiment. Agarose gel was utilized as it allowed for the creation of more accurate tissue-like scattering media directly around the vessel. Nonpolar polymers, such polydimethylsiloxane (PDMS), cannot mix with polystyrene bead solutions, making it difficult to control the optical properties of such polymers. Controlling the optical properties of the surrounding media was important to properly evaluate the performance of SC-OCTA and OCTA because a large portion of the signal comes from the ‘shadowing effect’ directly below the vessel (Figure S6 (c)).

### SNR Calculations

The SNR was calculated by the following:

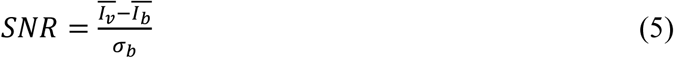
where 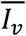 is the average vessel intensity, 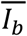 is the average background intensity, and *σ_b_* is the standard deviation of the background intensity. In the phantom measurements, the standard deviation of the SNR was calculated over 10 equally sized regions of interest (Figure S6 (f)). Statistical analysis of the phantom SNR was performed using a two-sample t-test.

### *In Vivo* Human Labial Mucosa Imaging

A healthy volunteer was recruited for *in vivo* labial mucosa imaging. The human lip was clamped down on a manually adjustable stage to allow the sample to be moved into focus (Figure S9). The subject was encouraged to only breath through their nose to prevent fogging of the objective.

### Sacrificed Mouse Imaging

Freshly sacrificed (< 2 hours postmortem) carcasses were carefully dissected and moved into focus using a 3D stage (Zaber, X-XYZ-LSQ150B-K0060-SQ3) (Figure S10). A c56BL/6 adult male mouse was used to image the outer surface of the ascending colon. An ICR (CD-1) adult female mouse was used for heart and anterior abdominal wall imaging. The mice were raised and sacrificed in accordance with Northwestern University IACUC standards.

## Supporting information

## Acknowledgements

We thank the O’Halloran Group, Dr. Wenan Qiang, and the Center for Comparative Medicine at Northwestern University for donating mouse carcasses for imaging and Rongrong Liu for providing knowledge on traditional OCTA. Additionally, we would like to thank Professor Oliver Cossairt at Northwestern University for suggesting Fourier ring correlation analysis. Support for this work was provided by the National Science Foundation Graduate Research Fellowship and the National Institute of Health (R01CA200064, R01CA183101, R01CA173745, R01CA165309). An award of computer time was provided by the INCITE program. This research used resources of the Argonne Leadership Computing Facility, which is a DOE Office of Science User Facility supported under Contract DE-AC02–06CH11357.

## Author Contributions

J.A.W. conceived the project, built the optical imaging system, and developed the spectral contrast codes. J.A.W. and G.S. acquired the data. L.M.A. dissected the mice and aided in interpreting the images. A.E. developed the edge detection code, produced spectra of different tissue types in Figure S7, computed the Fourier ring analysis, and developed the SC-OCTA simulations. J.A.W., A.E., and G.S wrote the manuscript, and L.M.A., T.Q.N, and V.B. aided in editing the manuscript and directing the project.

