## Supplementary material for "Spectral Contrast Optical Coherence Tomography Angiography Enables Single-Scan Vessel Imaging"

**Title**

### Supplementary Information

### Supplementary Note 1 – Visible OCT Oximetry & Angiography

There are several bodies of work that utilize visible spectroscopic optical coherence tomography (OCT) to analyze the spectral shape of hemoglobin and obtain microvasculature information. By assuming that the absorption spectrum of blood consists of a linear combination of oxygenated and deoxygenated hemoglobin, visible spectroscopic OCT has been used to map vasculature and quantify hemoglobin concentration within live retinal vessels[1]. However, this analysis required repeated scanning of the target volume 100 times and performed angiography through first high pass filtering of the temporal structural signal and then fitting to hemoglobin spectra. The work from this group was later improved upon by detecting the lower vessel location on the structural OCT image to better determine vessel oxygenation and only repetitively scanned the sample three times[2]. Other groups showed that one could perform spectroscopic angiography through a phase-sensitive decorrelation algorithm (similar to the one used for OCTA in this letter) and obtain vessel oxygen saturation with only five repetitive scans in the eye and brain, approaching capillary level vessels[3,4]. Yet, all the aforementioned techniques used repetitive scanning to locate vessels and therefore are highly sensitive to sample motion and require flowing blood. One body of work presents a capillary-level sensitive single scan angiography approach by imaging vessel shadows projected onto the outer retina using visible OCT[5]. This work used an *en face* slice of the outer retina to detect the vessel shadows and displayed the inverse OCT image intensity to produced two-dimensional vessel maps. However, unlike SC-OCTA, this approach would be sensitive to any highly attenuating structures, not just blood, and does not provide three-dimensional imaging of blood.

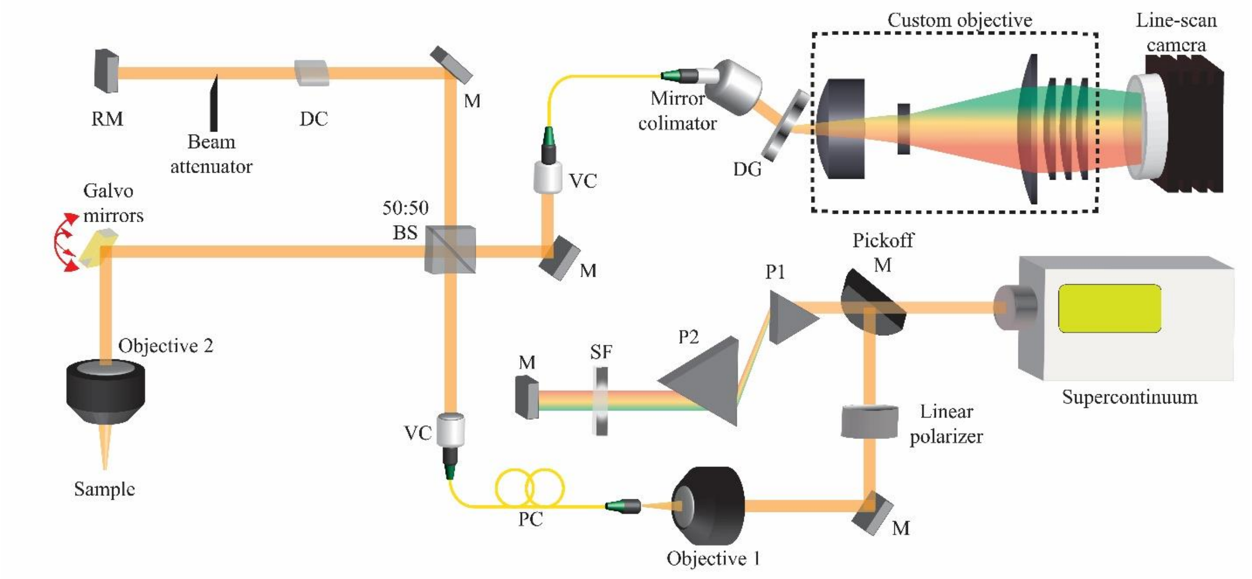


**Figure S1.** OCT System Schematic. Prism (P1 & P2). Spatial Filter (SF). Mirror (M). Paddle Polarization controller (PC). Visible collimator (VC). Beam splitter (BS). DC (Dispersion Compensator). Reference mirror (RM). Diffraction grating (DG).

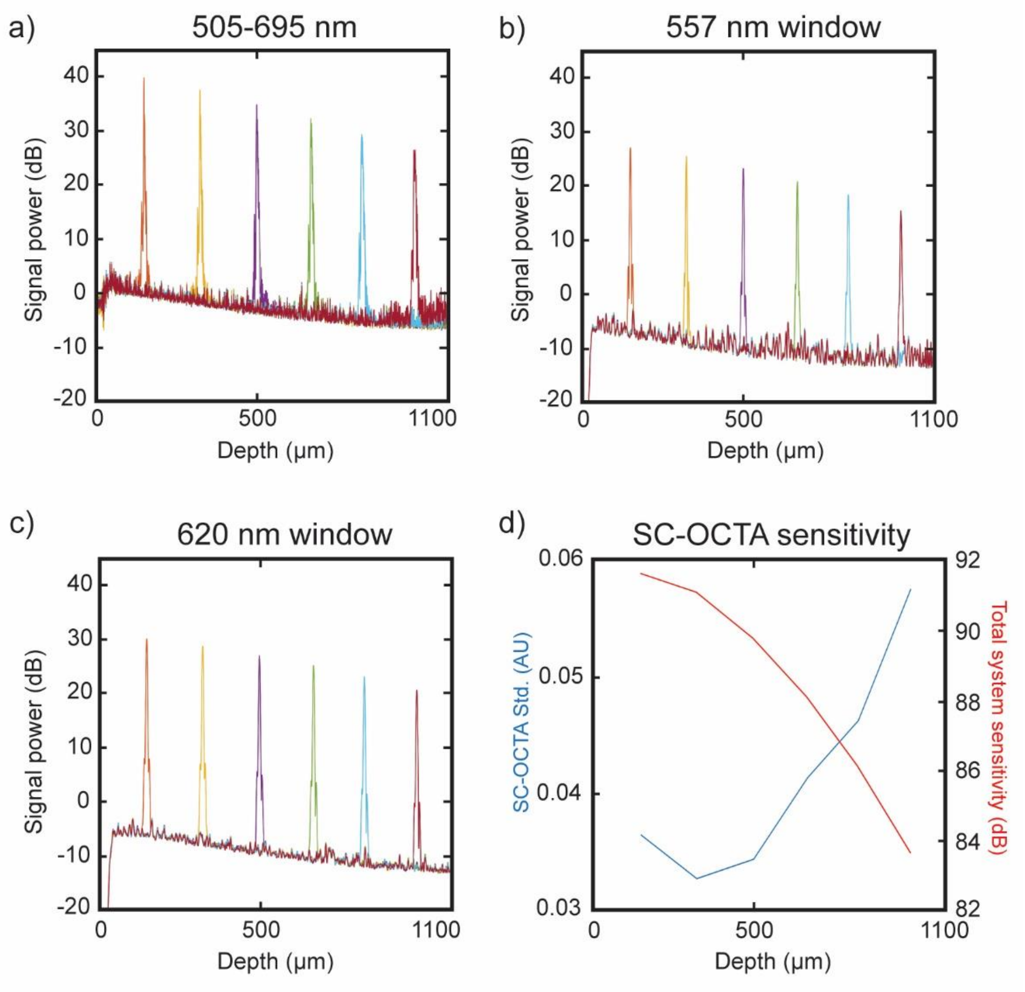


**Figure S2** Systems sensitivity measurements. The impulse response function of a mirror placed in the sample arm. The reference mirror position was changed to record the impulse response functions at different path lengths (depths) in air. These measurements should not be confused with system performance into different depths of a tissue sample, which has the added elements of sample optical attenuation properties and the axial point spread function of the focusing objective. Measurements were the average over 500 A-lines with a total round-trip attenuation in the sample arm of 51.4 dB using a neutral density filter. (a) Roll-off impulse response for the total system 505-695 nm bandwidth. Roll-off sensitivity ~-10 dB/mm. Air axial resolution (1.53 μm) and sensitivity (91.61 dB) measured from the first peak. (b) Roll-off impulse response for the 557 nm centered Kaiser sampling window. Roll-off sensitivity ~-14 dB/mm. Air axial resolution (3.80 μm) and sensitivity (86.05 dB) measured from the first peak. (c) Roll-off impulse response for the 620 nm centered Kaiser sampling window. Roll-off sensitivity ~-11 dB/mm. Air axial resolution (4.72 μm) and sensitivity (81.11 dB) measured from the first peak. (d) SC-OCTA signal standard deviation (blue line) and total system sensitivity (red line) measured in (a). The standard deviation of SC-OCTA signal was processed according to equation (3) for each mirror position over 500 A-lines. The aqueous 80 nm bead calibration was not necessary for estimation of standard deviation of SC-OCTA, and no median filters were used. The correlation between increasing system sensitivity and decreasing SC-OCTA standard deviation can be seen. We hypothesis the slight increase in standard deviation of SC-OCTA near the zero depth is a result of direct current noise which we attempt to minimize through high pass filtering the interferogram.

**Supplementary Note 2 – Theoretical Estimation of SC-OCTA SNR**

A macroscopic simulation based on the single scattering response was developed to numerically consider the contrast limits of the SC-OCTA signal with tissue noise and our system noise. We consider the OCT backscattered intensity can simplified with the following analytical expression:

$$I^2(x,y,z,k) = r\ L\ I_0^2(x,y,z,k)\ \frac{\mu_b(x,y,z,k)}{4\pi} exp(-2\ \mu_t(x,y,z,k)) \qquad \text{(S1)}$$

where $r$ is the reflectance of the reference arm, $L$ is the temporal coherence length of the source, $I_0$ is the incoming beam intensity, $\mu_b$ is the backscattering spectra, $\mu_t$ is the attenuation coefficient, *k* is the wavenumber in free space and *z* is the depth position in the sample. First, the geometry of the sample was established; for this simulation, we considered a cylindrical tube of blood with varying diameters, positioned 70μm below the surface and embedded in tissue. Each A-line is separated into its homogenous regions (tissue/blood/tissue) and the above expression was evaluated for each region, with its incoming intensity ($I_0$) modulated by the media above it, eg. reflecting with backscattered spectra at the interface and decaying within homogenous regions according to equation S1. This was repeated until the entire volumetric scattering intensity $I(x,y,z,k)$ was computed. The backscattering coefficient of tissue was assumed to have a power law $k^{(4\text{-D})}$ relationship, using a $D$ of 2.1. The absolute backscattering spectrum at each interface (tissue/air and tissue/blood) was normalized to yield a mean Fresnel reflection coefficient between their respective boundaries. The attenuation coefficient of tissue was taken from published healthy colon mucosal tissue[6], and the attenuation coefficient of whole oxygenated blood was taken from literature averaged values[7]. We then consider that the optical properties of a single RBC can be approximated by a volume equivalent bead of 3$\mu$m with refractive index matching that of oxygenated hemoglobin and background of tissue (refractive index = 1.38), and computed by Mie theory[8].

Then we consider the structural variation of tissue. We assume it is constant in wavelength, normally distributed, and can be represented as a constant scaling of the backscattered intensity. We computed the structural variation of the OCT image for labial mucosal tissue over 440 x 440 x 150μm area after a log transform. The mean normalized distribution had a standard deviation of 0.7; eg. Standard Deviation[log(Img3D)] / Mean[log(Img3D)]. A normally distributed random variable was added to the intensity I(x,y,z,k) with said standard deviation. The SC-OCTA signal was then generated according to (3). Next, we add the system noise to the SC-OCTA signal, which we quantified in Figure S2 (d), as a Gaussian distribution with a standard deviation of 0.03. This allowed for simulated SC-OCTA B-scans to be generated. An example of a simulated SC-OCTA B-scan with a 20 $\mu$m diameter vessel in tissue can be seen in Figure S3 (a).

From the simulated SC-OCTA B-scans we generated *en face* projections over 140 $\mu$m in depth and quantified the *en face* vessel line profiles which are shown in Figure S3 (b) for various sizes.

The $3\mu$m and $4\mu$m were generated using Mie theory, while the 10-50 $\mu$m diameter vessels assumed the optical response of whole blood. Furthermore, we include experimentally measured line profiles (L1-L3) from Figure 1 (e) and Figure S5 (a). The line intensities of L1-L3 were fitted with a Gaussian curve to measure the FWHM, which is listed in the legend of Figure S3(b).

While this simulation has many limitations, we see simply based on the contrast of $\mu_b$ and $\mu_t$, that SC-OCTA technique could show contrast for a single cell assuming the properties of a $4\mu$m bead and thus single capillary depending on the refractive index contrast, surrounding tissue fluctuations, and depth of integration. The floor of the line profile is characterized by the slope of the background tissue (eg. smaller $D$, will result in a steeper decaying $\mu_b$ and more negative background SC-OCTA signal). Thus, the optimal threshold for distinguishing vessel from tissue would need to be optimized based on the tissue type. Furthermore, SNR is sensitive to the homogeneity of the tissue and the refractive index contrast between the blood and tissue. SC-OCTA contrast will be greatest in more weakly scattering tissue. For example, if the background tissue refractive index is changed from 1.38 to 1.33, the *en face* 4 μm bead intensity doubles. Finally, it should be noted, that like all OCT imaging techniques, vessel response will decrease with depth into the sample due to system sensitivity roll-off, focusing and sample light attenuation.

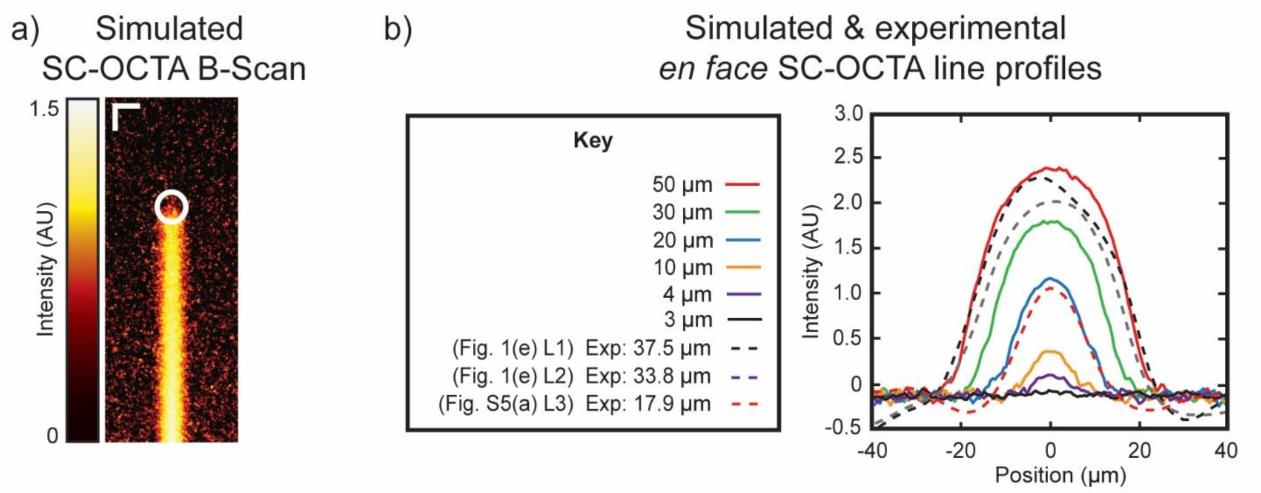


**Figure S3.** Macroscopic SC-OCTA simulation. (a) simulated SC-OCTA B-scan image for a 20 μm diameter vessel placed at the white circle. Scale bars: 20 μm. (b) Line profiles of simulated (solid) and experimental (dotted) *en face* SC-OCTA images integrated over 140 μm in depth. The experimental line profiles come from the SC-OCTA *en face* projections shown in Figure 1 (e) and Figure S5 (a).

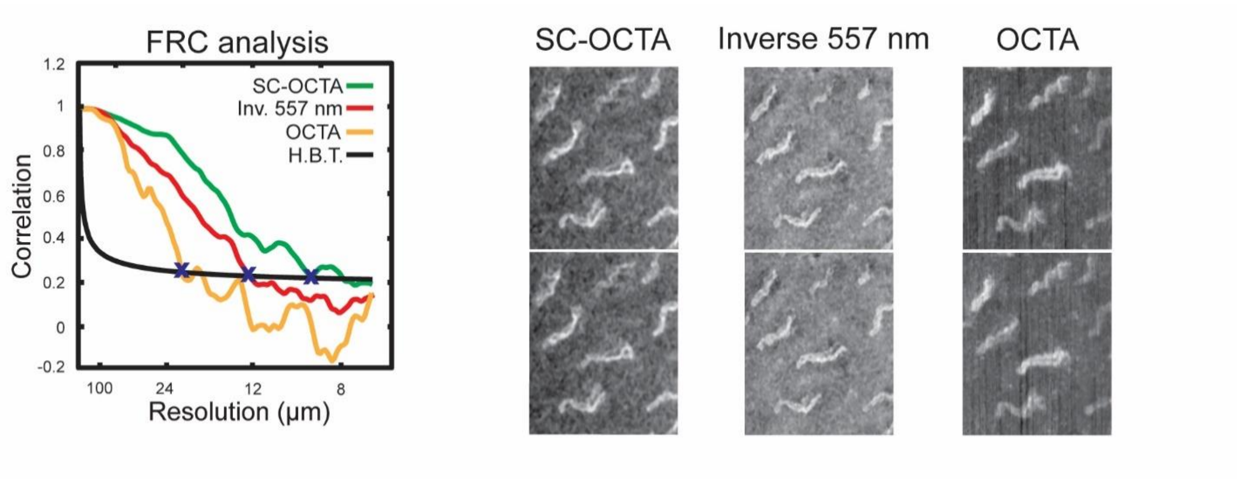


**Figure S4** Fourier Ring Correlation of the field of view in Figure 1 (c) for SC-OCTA, inverse 557 nm, and traditional OCTA. H.B.T (Half-bit threshold). As described previously[9]**,** the Fourier Ring Correlation (FRC) allows one to determine which feature sizes or spatial frequencies are conserved between subsequent measurements and which are attributed to noise by computing the Fourier correlation between successive data sets. For this analysis two images from OCTA, inverse 557 nm, and SC-OCTA were generated. For OCTA we used the angiography signal produced from two subsequent frames (frame 2 and 3 for one image, and frame 3 and 4 for another) not four subsequent frames like the rest of the OCTA images shown in this letter. For the inverse 556 nm and SC-OCTA images we used repetitions 2 and 4. Each pair of images were processed using the published FRC algorithm[9], and the crossing of the FRC curves with the half-bit threshold (blue 'x' on graph) was used to calculate the effective resolution of 20.19, 12.20, and 8.92 μm for the traditional, inverse 557 nm and SC-OCTA methods, respectively. Since the absolute diameters of the underlying vessels in that region were unknown, this metric can only implicate the relative effective frequency responses of each technique in an *in vivo* imaging scenario.

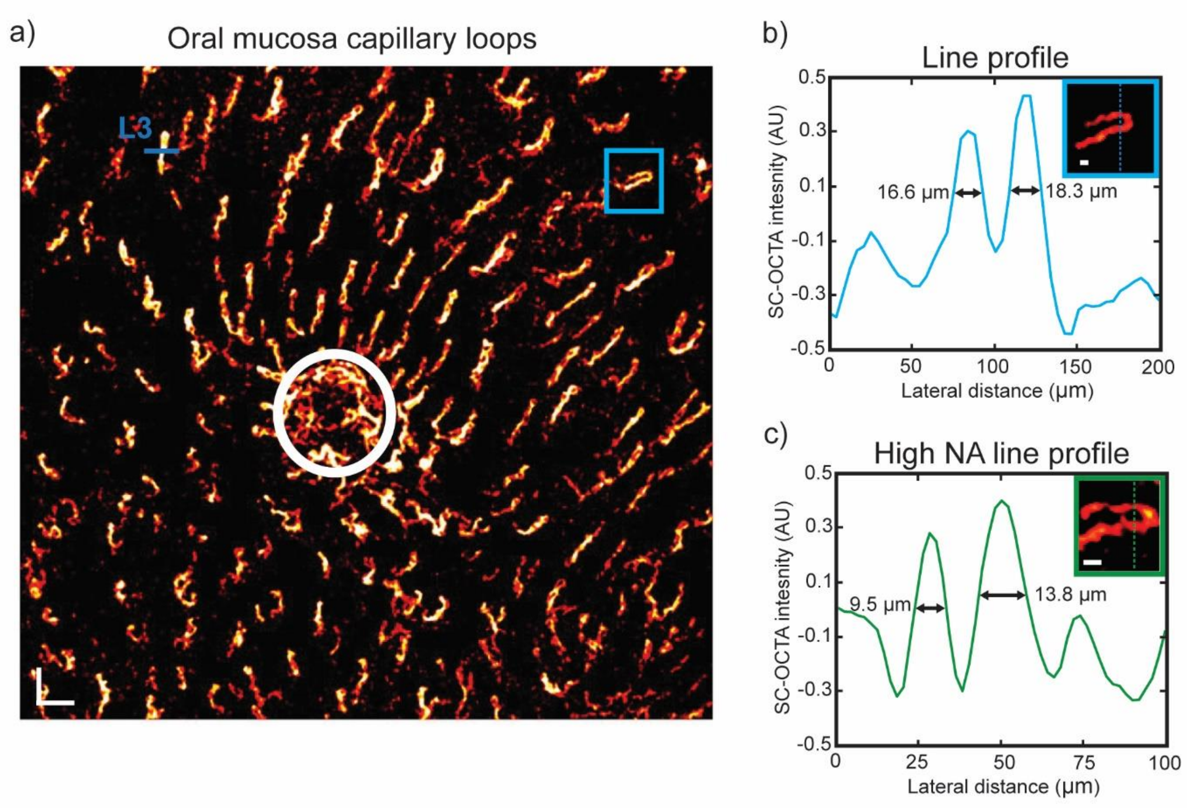


**Figure S5** (a) SC-OCTA *en face* projection (Depth: 70-209 µm) of capillary loops in labial mucosa of field of view (FOV) shown in Figure 1 (d & e). Capillary loop density for area outside of salivary duct (white circle) was calculated to be 14.65 loops/mm$^2$ which falls within reasonable physiological range according to a previous study[10]. Line 3 (L3) is line profile in Figure S3 for comparison with simulated line profile results. Scale bar: 200 µm. (b) Line Profile for the capillary loop shown in the blue box in (a) with FWHM for each peak across the loop. Scale bar: 20 µm. (c) To more accurately measure vessel diameter a beam expander was placed in the sample and reference arm (to account for dispersion) to increase the effective numerical aperture (NA) of the sample focusing objective resulting in a new lateral resolution of 2.1 µm. A 1.76 mm x 1.76 mm scan was taken with a sampling density of 2 µm. A capillary loop was then imaged over a new FOV with the higher effective numerical aperture in the lip. Please note this is not the same capillary loop shown in (b). Scale Bar: 20 µm. FWHM was calculated by fitting a Gaussian to each peak across the loop.

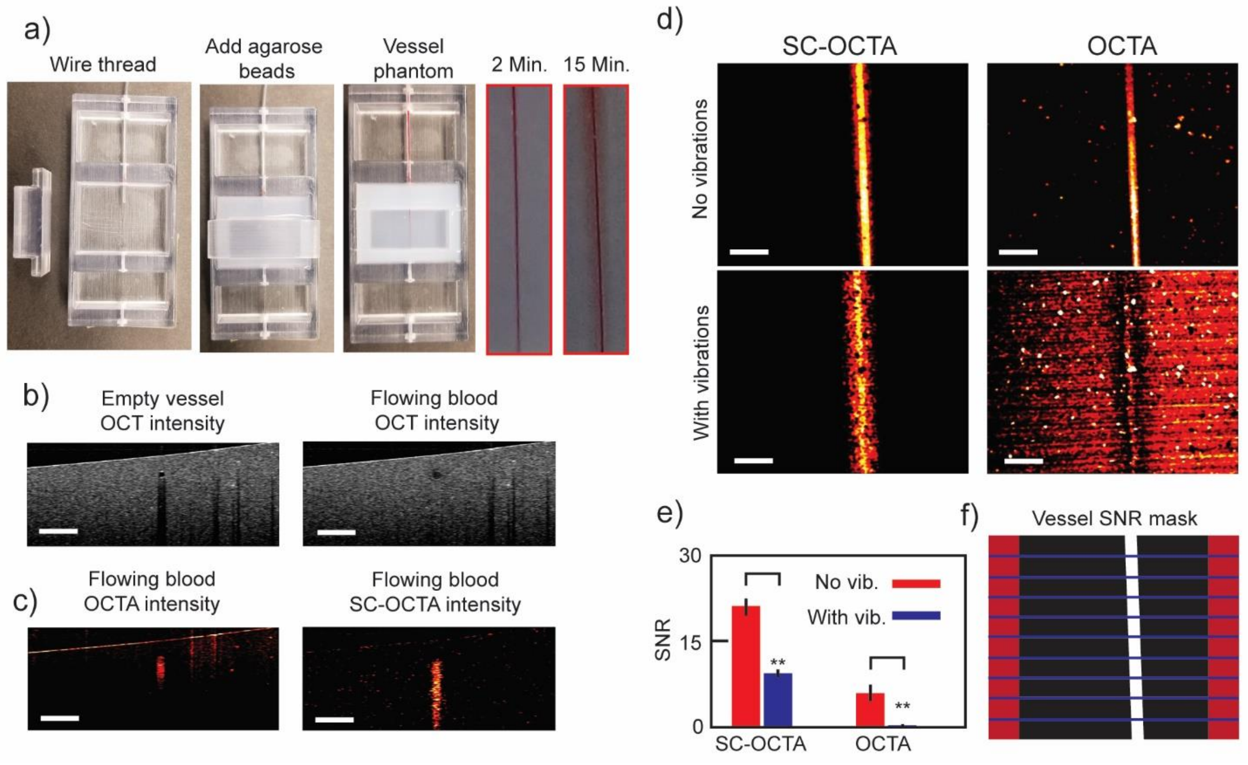


**Figure S6** (a) Construction of blood vessel phantom and photographs taken with smart phone of blood diffusion being observed at 2 and 15 minutes after flowing blood through the phantom. (b) OCT Intensity B-scans of vessel phantom before and after flowing blood through. From OCT image diameter was measured to be ~45 µm before blood was flown through and ~55 µm after, with a syringe pump set to .0006 µL/sec. A shadow is cast beneath the empty vessel because of reflections between air and the vessel wall. (c) Cross sectional examples of OCTA and SC-OCTA of vessel phantom with flowing blood. (d) SC-OCTA and OCTA *en face* projection with flow and flow with vibrations. (e) SNR analysis of *en face* projections in (d). SC-OCTA (No vibrations: 21.04±1.51 ; With vibrations: 9.34±0.65). OCTA (No vibrations: 5.904±1.57; With vibrations: 0.18±0.2). ** ($p<.01$) for two sample t-test. (f) Mask used for calculating SNR. Red is background. White is vessel. Blue lines delineate the 10 areas for determining standard deviation of SNR. All scale bars: 200 µm.

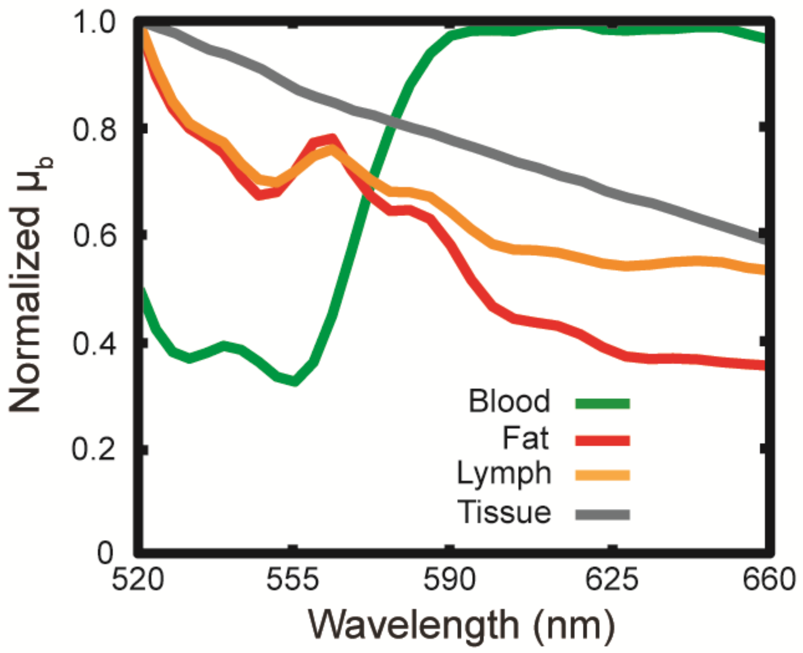


**Figure S7** Plots of normalized median backscattering ($\mu_b$) spectra from fat, lymphatic, blood vessels, and tissue measured by visible OCT in Figure 4 (a-d). Tissue backscattering spectra showed a gradual decay with increasing wavelength which has been previously noted[11]. Adipose and lymphatic tissue backscattering spectra decreased rapidly, behaving like Rayleigh scatterers. Their weakly scattering nature allowed them to be easily distinguished from higher scattering tissue with the inverse 557 nm spectral window. Blood had a characteristic absorption peak at ~550 nm, which was sensed by SC-OCTA. The spectra from adipocytes and lymphatic vessels are similar, which is expected since lymphatic vessels are associated with fat transport in abdominal areas[12].

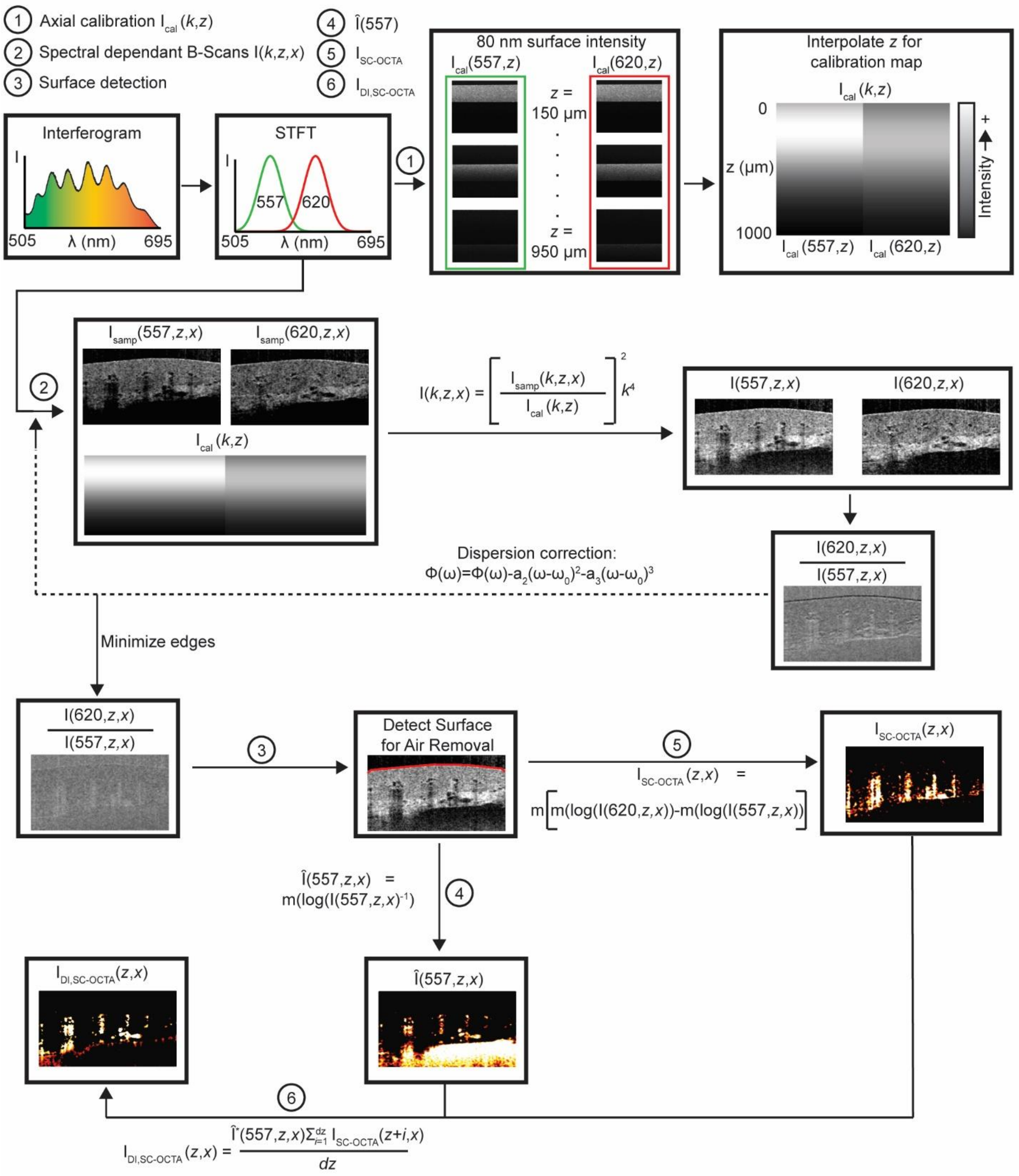


**Figure S8** Flow chart showing processing steps for generating spectral-contrast-based angiography images ($\hat{I}(557)$, $I_{SC-OCTA}$, $I_{DI,SC-OCTA}$). The coordinate $z$ refers to the depth direction along an A-line and $x$ is the b-scan direction. m is referring to a 3D median filter that is applied to the $z$, $x$, and C-scan direction.

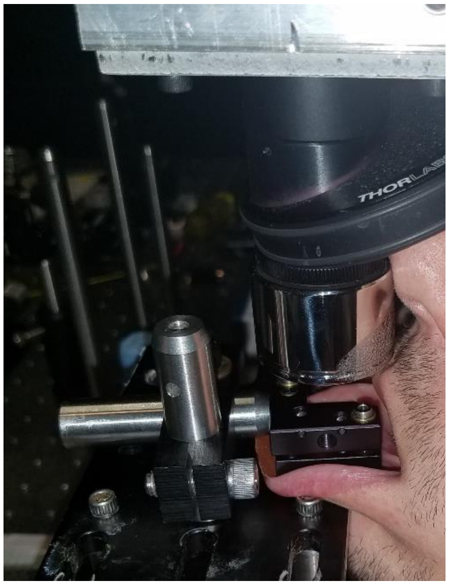

**Figure S9** Procedure of *in vivo* human imaging of labial mucosa. Lip was clamped down and sandpaper is used on clamp to prevent slippage. A hand-driven stage allows for fine adjustment of lip location.

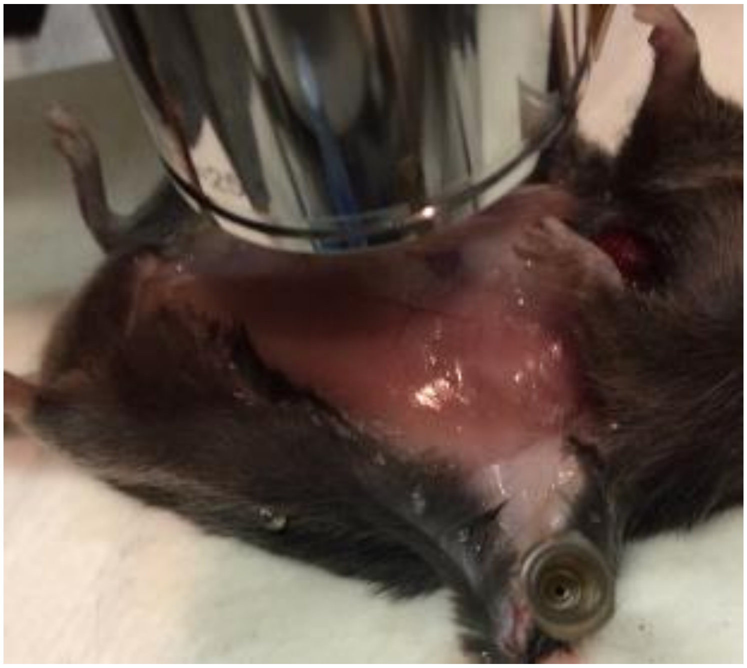

**Figure S10** Imaging of freshly sacrificed mouse tissue.

### Videos

**Video S1** 3D rendering of FOV from Figure 1 (d & e). Full spectrum OCT intensity (gray), inverse 557 nm (white/orange), and Depth Integrated SC-OCTA (Green).

**Video S2** 3D rendering of freshly sacrificed mouse heart. Full Spectrum OCT intensity (gray) and Depth Integrated SC-OCTA (green). Note this is not the same field of view shown in Figure 4 (e).

**Video S3** 3D rendering of the lymphatic fluid around a lymphatic valve from LV1 in Figure 4 (d).

**Video S4** 3D rendering of FOV from Figure 4 (c). Full Spectrum OCT intensity (gray), inverse 557 nm with blood vessels removed (white/orange), and Depth Integrated SC-OCTA (green).

**Video S5** B-scans of vessels phantom during vibrations.
